# Data-driven streamflow modelling in ungauged basins: regionalizing random forest (RF) models

**DOI:** 10.1101/2020.11.14.382598

**Authors:** Arnan Araza, Lars Hein, Confidence Duku, Maurice Andres Rawlins, Richard Lomboy

## Abstract

Streamflow predictions in ungauged basins (PUB) has been geared towards data-driven methods, including the use of machine learning methods such as random forest (RF). Such methods are applied in PUB regionalization or the transfer of a streamflow model from gauged to ungauged (sub) basins or watersheds after grouping watersheds on similarity rules. Regionalized streamflow models are needed for tropical-mountainous regions like Luzon, Philippines - where gauged data is limited - but demands on streamflow modelling for water resources accounting and management are high. In 21 watersheds in Luzon, we “regionalize” RF streamflow models after grouping watersheds based on: a principal components analysis (PCA-clustered), by major river basins (basin-clustered), whole study area scale (one-clustered). Another method without watershed grouping (watershed-level) was also included and inter-compared. Among the four methods, goodness-of-fit evaluations revealed that PCA-clustered method was higher by at most 0.35 coefficient of determination *R*^2^ and 0.31 nash-sutcliffe efficiency *NSE*, and the least bias in 8 of 12 monthly flows. These are attributed to the added-value of homogeneous watershed grouping, reflected by higher importance (to RF models), of static covariates from open and high-resolution data. Normalized errors from monthly streamflow showed a clear bias (least with the PCA-clustered metohd), linked to season and water management practices in the study area. Ungauged watersheds in the Philippines can effectively use streamflow models from gauged watersheds if they belong to the same cluster.

**Key Points:**

- Ungauged sub-basins (watersheds) can effectively use RF streamflow models from gauged watersheds if both belong to the same cluster.
- Biophysical data from high-resolution open data are valuable: as basis for watershed clustering and as streamflow predictors to RF models.
- The predicted streamflow from the RF models reflects seasonal deviations in streamflow caused by natural and man-made water regulation

## 1 Introduction

Globally, water resources are increasingly under pressure from over-exploitation, deforestation affecting streamflow patterns, and climate change (Oki, 2006). Better managing river water resources requires detailed information on streamflow and how streamflow depends upon river basin management. However, in many tropical and developing countries, in-situ streamflow monitoring networks are neither adequately maintained nor have sufficient spatial coverage in the river basins (Do et al., 2018). This is also the case in the Philippines where streamflow data scarcity and paucity have hampered credible streamflow estimates for major river basin management (Tuddao Jr, 2009).

Streamflow prediction in ungauged and poorly gauged basins (PUB) is commonly performed using hydrological model-based approaches (Hrachowitz et al., 2013). These include process-based methods which depend on physics-based natural processes. Process-based methods are parametric, which make use of model and input-specific parameters to perform a water balance simulation using empirical equations (Arnold et al., 2012). Process-based models are often implemented in user interfaces, enabling input-level modifications to model not only business as usual events, but also land-use change, climate change, drought, and land management scenarios in sub-basins (herein referred as watersheds) scales (Jaranilla-Sanchez et al., 2011; Mango et al., 2011; Palao et al., 2013). For large scale-applications, e.g. in multiple states (Mohamoud, 2008) or at pan-European scale (Pagliero et al., 2019), watershed regionalization is often applied first before executing a hydrologic model. Regionalization includes transferring hydrological model and(or) model parameters from gauged watersheds to ungauged ones in a “donor-receiver” logic. Transfer is either based on watershed-to-watershed distance (like spatial proximity) or based on regression of hydrological parameters and watershed characteristics (Beck et al., 2016). A review by Razavi & Coulibaly (2013) identified at least 30 unique regionalization studies conducted in the past decades mostly for engineering applications. On the other hand, they also categorized PUB as hydrological model-independent or a non-parametric approach that bypasses hydrological models and its parameters. In this case, the watershed characteristics functioning as explanatory variables of a gauged watershed are calibrated directly with streamflow itself to create a “streamflow model” usable to ungauged watersheds. This empirical method has emerged as an alternative to model-dependent methods for practical reasons of non-parameterization and faster implementation (Loukas & Vasiliades, 2014). Moreover, conducive to this approach is the advent of input-level big data, redefining model-independent as data-driven methods (Razavi & Coulibaly, 2013). Contextually, a data-driven method is defined as the empirical and non-physics-based calculations in river basins using considerable amount of data and machine learning (Solomatine & Ostfeld, 2008).

The broad state-of-the-art application of machine learning (ML) in water resources modelling has been categorized to forecasting, preprocessing, variable selection, training-testing dataset optimization, and predictive performance assessments (Tyralis et al., 2019). One of the most used ML methods in water resources is Random Forest (RF), a supervised algorithm for predictive modelling developed by (Breiman, 2001). Currently, RF is widely-used in solving regression-based problems and is perceived to solve more complex water resources problems in the future (Tyralis et al., 2019). When applied to streamflow modelling, comparatively, RF has shown more optimal streamflow predictions (Papacharalampous & Tyralis, 2018) and slightly better streamflow model accuracy (Shortridge et al., 2016) than other ML methods. But what makes RF unique, aside from its predictive power and efficiency in non-linear relationships, is the capability to provide importance values to explanatory variables (Tyralis et al., 2019). For example, Bruner et al., (2018) showed that physical watershed parameters explain most of the RF streamflow model variability to predict flood hydrographs and (Shortridge et al., 2016) highlighted the sensitivity of streamflow models to temperature as proxy to climate change effects. Furthermore, uncertainties in RF model predictions are quantifiable (Wager et al., 2014). But in watershed regionalization, RF and so with other ML methods, are mostly used only as a complementing model in regionalizing parameters of hydrological models (Saadi et al., 2019; Brunner et al., 2018; Senent-Aparicio et al., 2019), large-scale but non-regionalized streamflow modelling (Asefa et al., 2006), and watershed classification or grouping (Yadav et al., 2007; Isik & Singh, 2008).

Pivotal to the success of regionalization is on how well watersheds are grouped or classified. While the choice of grouping method is subjective to basin characteristics and study area scale (Saadi et al., 2019), “homogeneous” classification of watersheds at regional-scale has been widely-tested. In similar studies, almost all of regional variability was captured when datasets are transformed into principal components, thus becoming the watershed grouping basis (Isik & Singh, 2008; Prieto et al., 2019). Such method was found effective to perform adequacy evaluation of streamflow models. In less heterogeneous regions, grouping neighboring watersheds together (e.g. inside a major river basin) is reasonable (Oudin et al., 2008). Effective grouping could facilitate the use of RF streamflow model itself to be “regionalized” in ungauged watersheds. For example, one performed watershed clustering and streamflow modelling to better predict monthly streamflow (Wu et al., 2010) while another did the same to predict flood indices (Latt et al., 2014). These examples both evaluated their models only at cluster level, and not at watershed-level e.g. by using pseudo-ungauged watersheds (Goswami et al., 2007; Athira et al., 2015). Moreover, streamflow models should be evaluated whether predicted streamflow shows systematic deviations (e.g. seasonal) or bias, but they are often disregarded (Arsenault & Brissette, 2014). Other similar studies confined regionalization of RF models at basin scales and none has attempted in tropical-mountainous regions.

Tropical-mountainous regions often have complex hydrological system with various natural and human-induced impacts on hydrological interactions (Wohl et al., 2012). Moreover, these regions have very limited streamflow data, but high streamflow modelling demands on water resources research and development (R&D) for better watershed management. Given these, the main objective of this paper is to test and compare how a regionalized, RF-based streamflow models can be used to predict streamflow in ungauged watersheds in mountainous regions of Luzon, Philippines. In line with that goal are the following research questions: (1) What is the effect of watershed grouping in regionalizing RF models? (2) How are open and high-resolution data useful in RF model regionalization? and (3) What influences seasonal bias in streamflow predictions?

## 2 Study area

Our study area includes 21 watersheds within six major river basins and covers five administrative regions in Luzon, Philippines. The study area is shown in Figure 1 where each watershed is labeled according to its respective major river basin and the river or station name e.g. crb_a is equivalent to Cagayan River Basin, Aligapay River.

**Figure 1.**
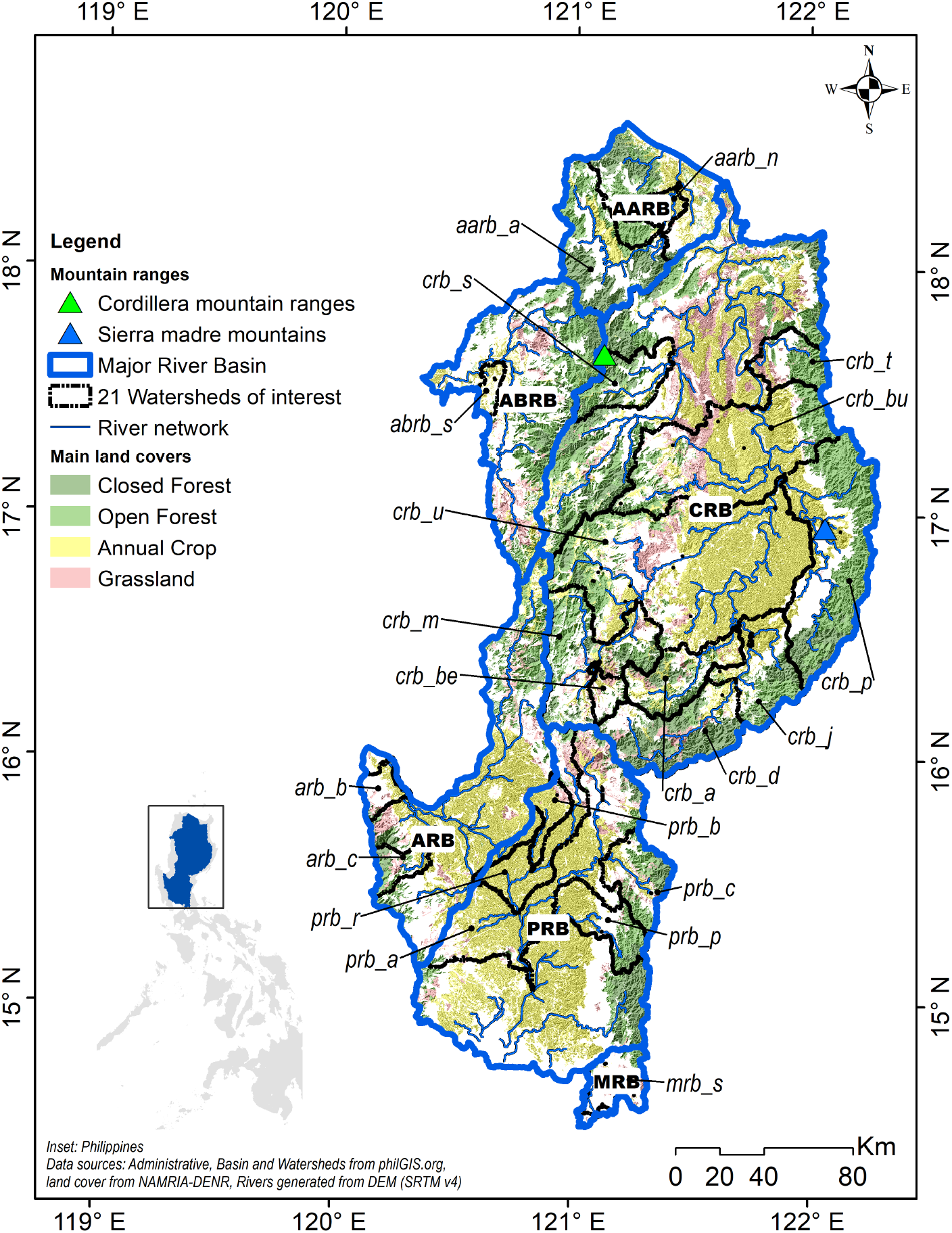
Study area overview showing the 21 watersheds within the 6 major river basins in Luzon namely Abra, Agno, Apayao-Abulog, Cagayan, Marikina, and Pampanga River Basins. The basins are labelled as *abrb, arb, aarb, crb, mrb*, and *prb*, respectively. River networks are shown also to guide upstream-downstream connections within watersheds while main land covers show how forested the uplands and cultivated the lowlands are.

Inside the study area are the top crop producing provinces of the country. This means that long-term water management for irrigation is very important to sustain crop production. The water for irrigation comes from the headwaters of *Sierra Madre* and *Cordillera* mountains - two of largest mountain ranges in the country. With mountain ridges up to 2000 meters high, the watersheds depict a ridge-to-floodplain form of irregular and elongated sizes ranging from 20 to 10,000 km^2^. This shape and size allow a sloping terrain topography with mostly mountain, silt, and clay soils. The forests in these area sustain river flows even in driest months of March and April. More than half of these forests (60%) are classified as public forest lands. The forests contribute to make more rain up to around 2000mm/year in most of the watersheds. In wet season, the lowland of the study area is a floodplain with high risks of flooding and landslide due to typhoons (Cinco et al., 2018). These impacts are aggravated by deforestation and climate change (Rawlins et al., 2016). As a response, the government formulated management plans in every major river basin focusing on long-term domestic water use, modernizing irrigation, and flood mitigation.

## 3 Methods

This study implements three stages to regionalize RF models into ungauged watersheds (Figure 2). The steps are elaborated in the succeeding sub-sections. As an overview, section 3.1 is about the open and high resolution data inputs turned streamflow predictors or covariates. Section 3.2 tackles the two-step nature of regionalization: watershed grouping and RF streamflow model “transfer”. Lastly, in section 3.3, predicted streamflow are evaluated relative to uncertainties from RF models and seasonal bias.

**Figure 2.**
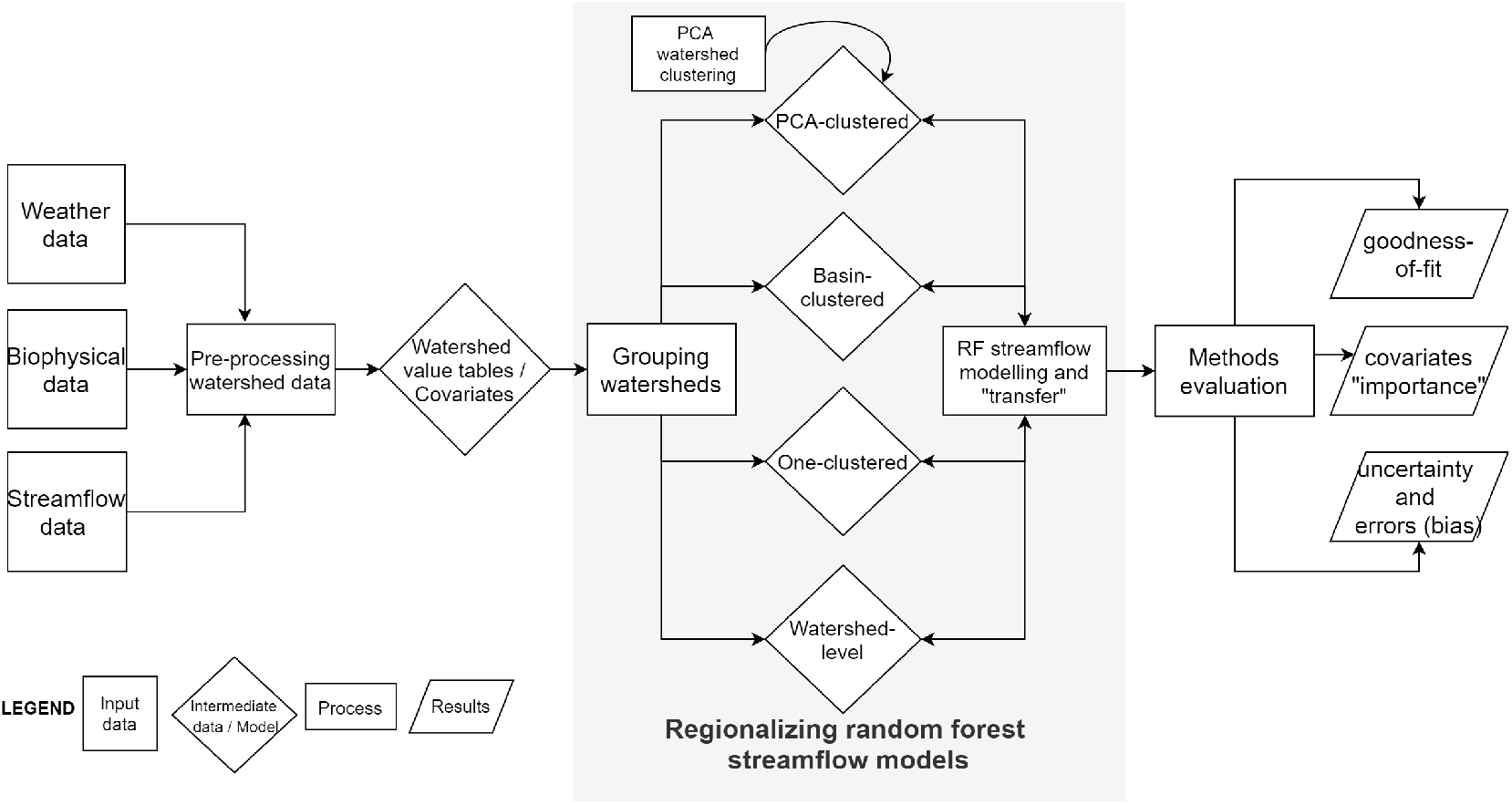
Overall diagram of the steps undertaken in this study starting from data collation, regionalization of RF models, and ending with models evaluation.

### 3.1 Input data and covariates

The basis of input data selection is to represent the main components of a tropical mountainous hydrological system including streamflow, biophysical, climatic, and other complementary data. Another basis is the ease of access from these data.

We obtained 51,690 records of daily streamflow data (herein referred as observed streamflow), ranging from 2000 to 2016, and expressed as *m*^3^/*s*. The observed data come from 21 gauged stations and computed by averaging three daily gauge measurement. Geo-tagging the stations allowed us to delineate 21 watersheds from an elevation raster using *ArcHydro* tool (Maidment & Morehouse, 2002). Within and nearby these watersheds, we obtained weather data from local monitoring stations. The gaps in daily weather data were interpolated using predictions from a linear regression model of the “gap station” and neighboring weather station. Vegetation inputs were represented by land cover data from 30m Global Land Cover (Arsanjani et al., 2016) and forest-loss from 30m Global Forest Change (Hansen et al., 2013). The latter was included to depict the forest loss history in the area. Soil inputs came from a global but high resolution soil dataset raster (250 m) from World Soil Information (Hengl et al., 2017). The dataset is a collection of model-based predictions using soil profile and environmental variables. Next, elevation was included to capture the ridge-to-floodplain form of the watersheds. For that, we used the 30m Shuttle Radar Thematic Mapper Digital Elevation Model Version 4 (SRTM-DEM V.4) (Jarvis et al., 2008), which has higher resolution and vertical accuracy than other similar options (Pryde et al., 2007). From the elevation data, slope was derived as the last main input. All rasters were projected into UTM Zone 51N then mosaicked and masked within the study area using Geospatial Abstraction Library (Warmerdam, 2008). After that, coarser rasters were resampled into 30m using nearest neighbor approach to match spatial resolutions. To better represent soil, elevation, and slope instead of using the mean, we reclassified them into five classes. We classified the rasters by quantiles to assure equal population within classes (with 5 as the highest value). As complementary watershed data, we included climate type, month, land area, and the major river basin where it belongs. The final pre-processing step was to overlay all inputs per watershed. All data pre-processing were implemented in *R* (R Core Team, 2013) in *R studio*.

All non-streamflow input data were used as covariates, also known as explanatory variables or predictors. From the main inputs, more covariates were derived and categorized as weather (W), physical (P), land cover (L), hydrologic (H), season (S) and complement (C). Based on daily rainfall data, weekly and monthly rainfall were derived to capture short-term and seasonal rainfall patterns. From the land cover, each land cover class was a separate covariate; and so were hydrologic parameters curve number and Manning’s roughness - these two are unique per land cover class. Similarly, each class from soil, slope, and elevation inputs was a stand-alone covariate. In total, 55 covariates were generated as shown in Table 1, where each main input has covariates count, label, measuring unit, and temporal unit. The labelling is intended not only for concise naming of covariates (e.g. for results), but also to emphasize the covariates category. The covariates are also categorized whether static or dynamic to assess how streamflow models are affected by temporal variations. Temporally, the covariates are balanced in number: weather covariates are the most dynamic with daily to monthly values, while the hydrologic covariates are yearly. A land cover class could be dynamic if forest conversion happens because of annual net forest loss. The climate and physical covariates are static for a reason since soil, slope, and elevation gradually change.

**Table 1.**
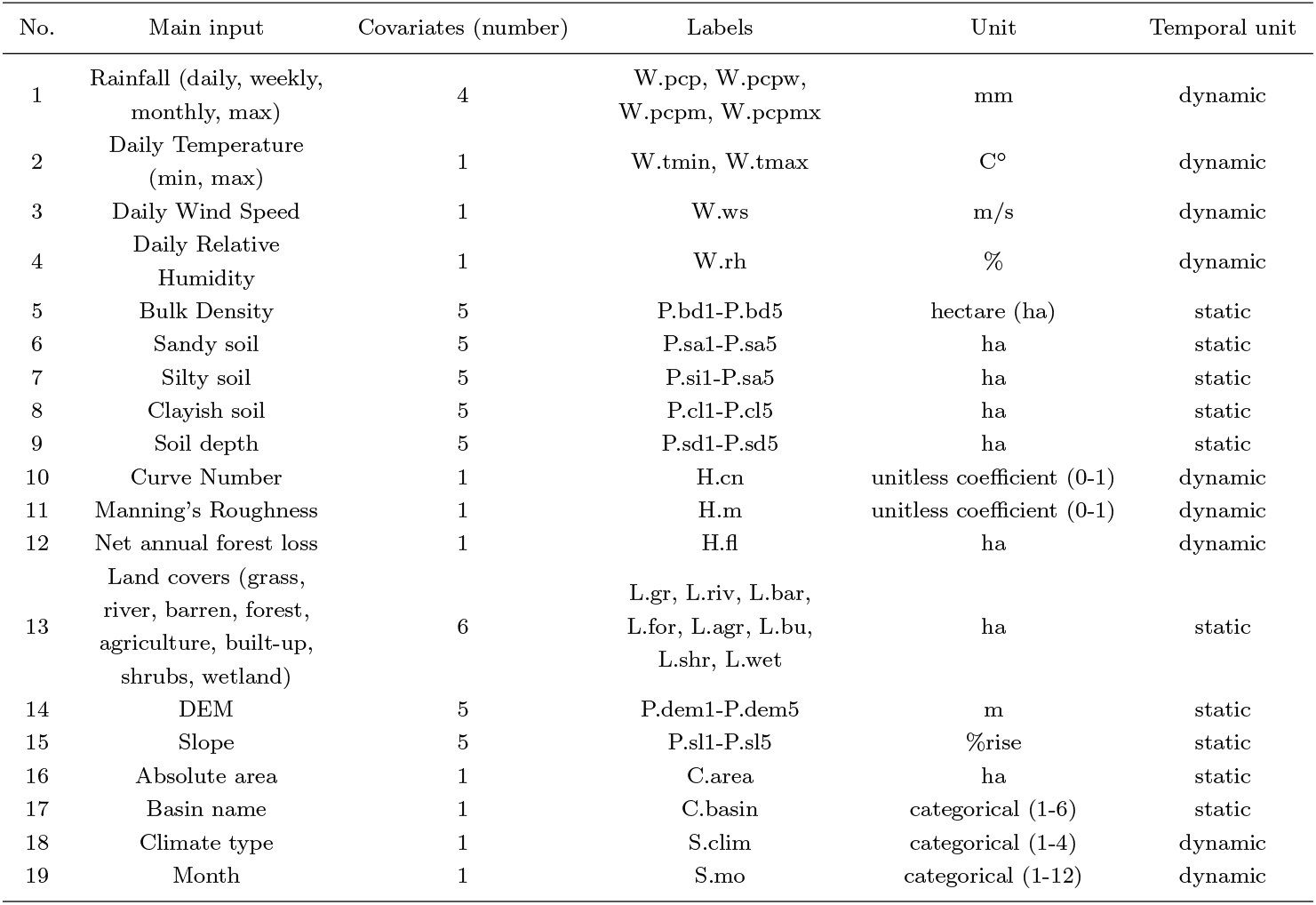
List and key information of main covariates including the main data inputs, number of covariates derived from main inputs, covariates labels, measuring unit, and temporal unit. Each watershed has an equivalent value table.

| No. | Main input | Covariates (number) | Labels | Unit | Temporal unit |
| --- | --- | --- | --- | --- | --- |
| 1 | Rainfall (daily, weekly, monthly, max) | 4 | W.pcp, W.pcpw, W.pcpm, W.pcpmx | mm | dynamic |
| 2 | Daily Temperature (min, max) | 1 | W.tmin, W.tmax | C° | dynamic |
| 3 | Daily Wind Speed | 1 | W.ws | m/s | dynamic |
| 4 | Daily Relative Humidity | 1 | W.rh | % | dynamic |
| 5 | Bulk Density | 5 | P.bd1-P.bd5 | hectare (ha) | static |
| 6 | Sandy soil | 5 | P.sa1-P.sa5 | ha | static |
| 7 | Silty soil | 5 | P.si1-P.sa5 | ha | static |
| 8 | Clayish soil | 5 | P.cl1-P.cl5 | ha | static |
| 9 | Soil depth | 5 | P.sd1-P.sd5 | ha | static |
| 10 | Curve Number | 1 | H.cn | unitless coefficient (0-1) | dynamic |
| 11 | Manning's Roughness | 1 | H.m | unitless coefficient (0-1) | dynamic |
| 12 | Net annual forest loss | 1 | H.fl | ha | dynamic |
| 13 | Land covers (grass, river, barren, forest, agriculture, built-up, shrubs, wetland) | 6 | L.gr, L.riv, L.bar, L.for, L.agr, L.bu, L.shr, L.wet | ha | static |
| 14 | DEM | 5 | P.dem1-P.dem5 | m | static |
| 15 | Slope | 5 | P.sl1-P.sl5 | %rise | static |
| 16 | Absolute area | 1 | C.area | ha | static |
| 17 | Basin name | 1 | C.basin | categorical (1-6) | static |
| 18 | Climate type | 1 | S.clim | categorical (1-4) | dynamic |
| 19 | Month | 1 | S.mo | categorical (1-12) | dynamic |

These covariates constitute the watershed information and were merged into a collated table we refer to as watershed value table. This table is unique per watershed and served as the formatted data for streamflow model training and evaluation using random forest.

### 3.2 Regionalizing random forests

As mentioned, RF has been widely applied into water resources research and development towards solving non-linear problems in hydrology through data-driven methods. That is in line with our expectations that RF can capture streamflow responses in a 16-year stretch that will allow RF streamflow models usable to ungauged watersheds. All that watershed information are placed in each watershed value table as input for regionalization.

#### 3.2.1 RF algorithm

Random forest algorithm by (Breiman, 2001) is a supervised learning model used for classification and regression. We used it for the latter because streamflow is a continuous variable. Conceptually, the algorithm works in five steps as demonstrated in Figure 3 in a single RF tree. First, a watershed value table is assembled, where observed streamflow and the covariates are placed in columns. Second, the watershed value table undergoes bootstrapping, where the rows are randomly selected (with replication) to form a sub-sample or simply a bootstrap sample of the watershed value table. Third, from this random subset, the root node (topmost circle of a tree, Figure 3) is randomly selected from a limited choices of covariates. The root node is split based on a yes-no decision - leading into two succeeding nodes - again coming from a limited set of covariates. Fourth, nodes increase (bagging or “tree growing”) on every binary decision until reaching the final decision. That bagging step de-correlates the covariates within the tree and assuring lesser prediction variance. That distinguishes RF from the unrestricted splitting of the traditional bagging method. Lastly, as trees grow in parallel, the final decision of each tree are averaged to get the final predicted streamflow. RF prediction can be written as:

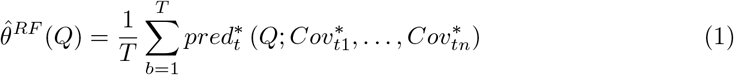

where streamflow is predicted 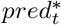 from an individual tree of an RF streamflow model 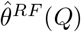 as the average prediction among the decision trees 1 to *B* with covariates 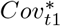 to 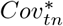.

**Figure 3.**
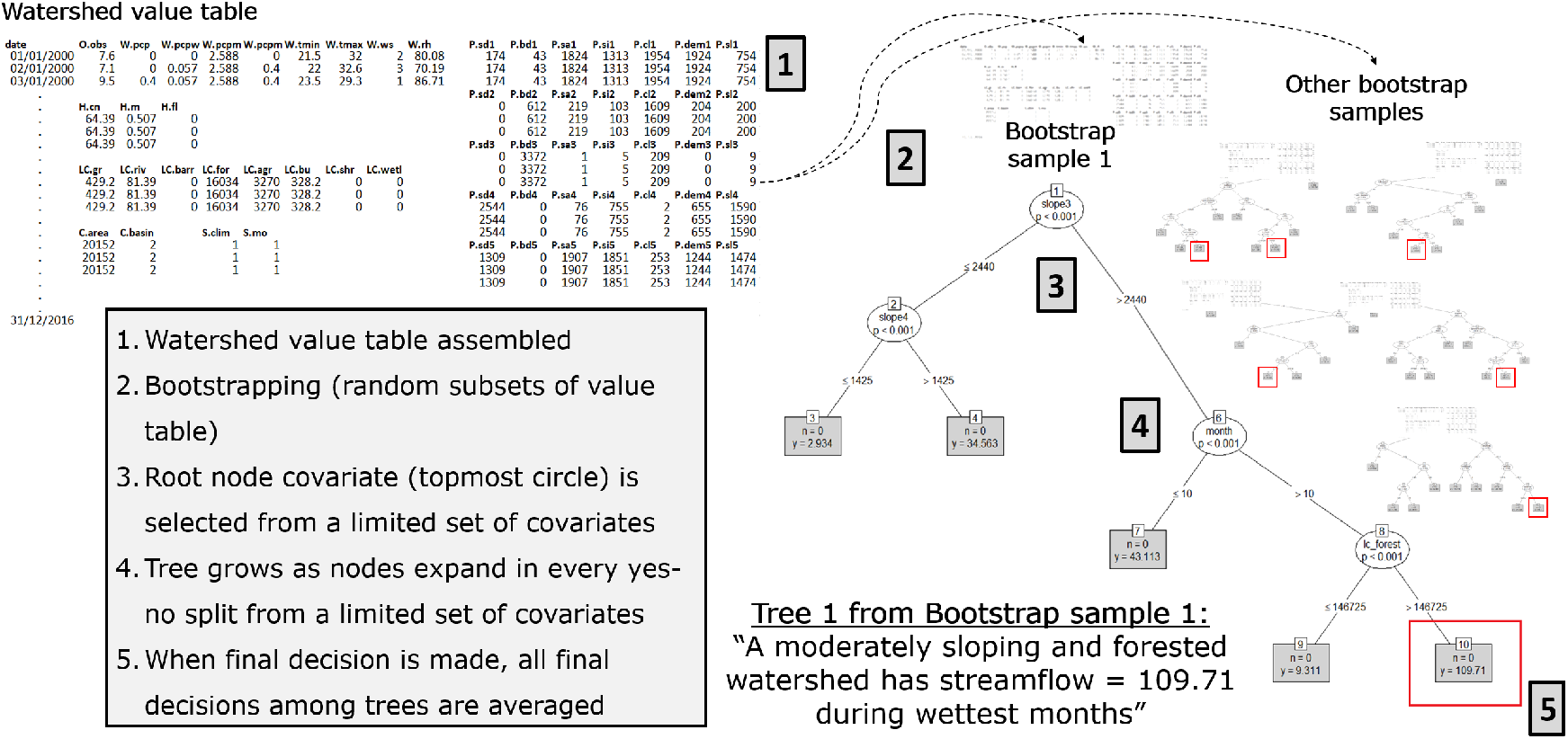
Summary of RF implementation highlighting one tree from a bootstrapped sample, to (restricted) tree growing, until the final predictions in red boxes. The quoted line in the diagram depicts the link of the predicted streamflow all throughout the root (topmost) node.

We used the RF algorithm using the *ranger* package in R for faster implementation (Wright & Ziegler, 2017). Prior to that, the RF hyperparameters were tuned to an optimal value. These parameters are: (1) the limited choices of covariates when trees grow (recall step 3, Figure 3), also called as the “split variable”; (2) another is the maximum number of bootstrap samples or trees to grow (“number of trees”) in a random forest. The two hyperparameters are termed as *mtry* and *num.trees* in *ranger* package). The correct value of the two considers the trade-off between overfitting and computation time. If the split variable is set too high, the model could overfit or see repetitive information, and produce highly correlated trees (Probst et al., 2019). Most likely, that will inaccurately predict streamflow from any new dataset. After the tuning, the split variable was set to be 1/4 of the total covariates while the default value of the number of trees (500) was retained.

#### 3.2.2 Regionalization methods

We define the term “regionalization” as the transfer of streamflow models from gauged to ungauged watersheds after a watershed grouping scheme based on similarity rules. The “grouping” not only assigns watersheds into groups, but also their information or watershed value tables for a certain RF model training. With this two-step approach, we trained RF models depending whether the watershed grouping is PCA-clustered, basin-clustered, and all-clustered. We also included a method without watershed grouping. These methods are depicted in Figure 4.

**Figure 4.**
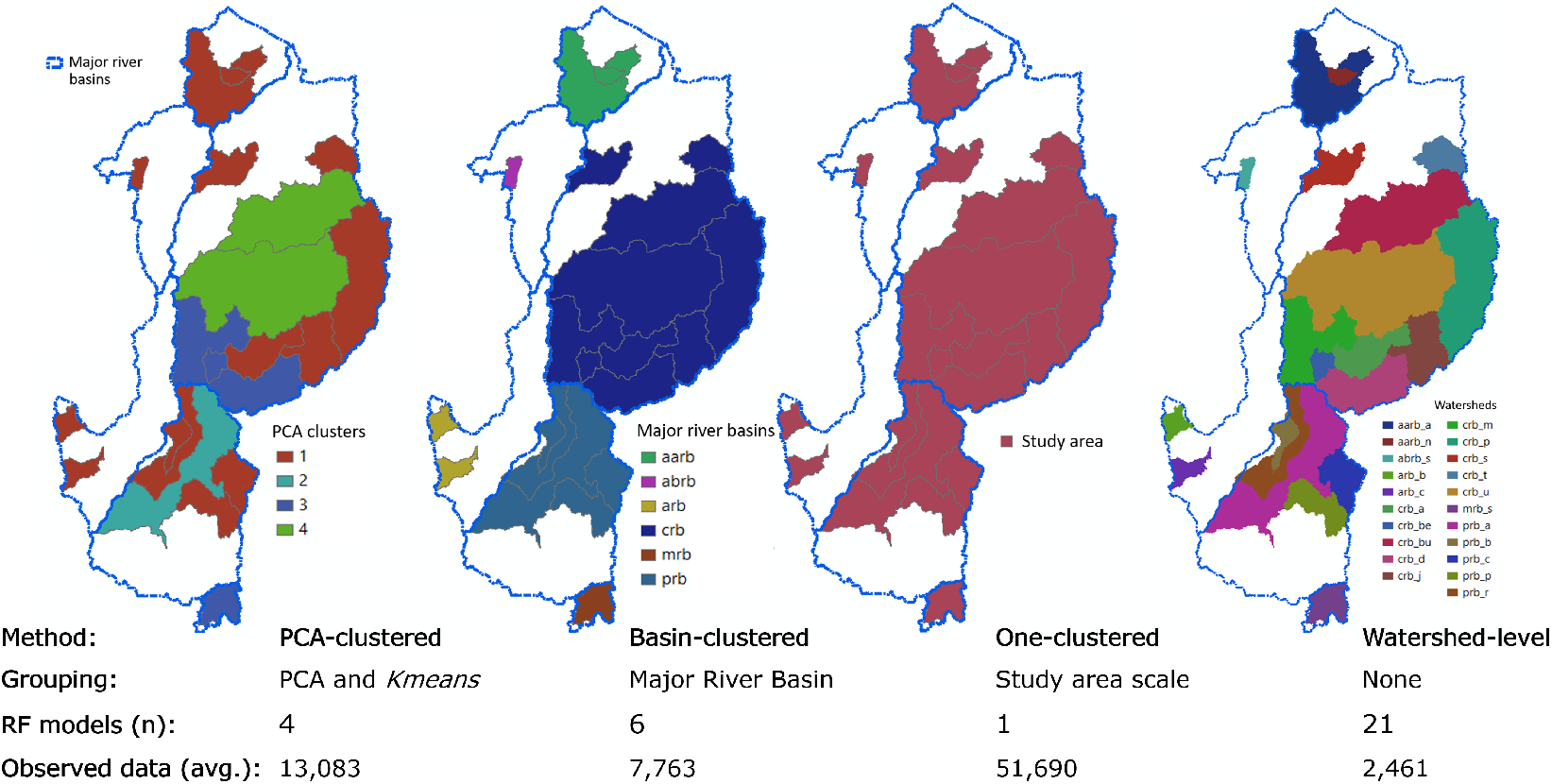
Maps of watershed grouping methods as basis for regionalizing RF models. Indicated on each method are the number of RF models and the average observed data per RF model.

The first method uses principal components analysis (PCA) to cluster watersheds into four, resulting to an RF model per cluster (elaborated next section). The second method assigns the watersheds to their mother river basin resulting to six RF models corresponding to each major river basin. This grouping method can signal whether basin-level variability is evident. The third method merges all watershed value table to train one RF model. In this case, we assessed if the RF model can still predict accurately from a training data with all watershed information at the whole study area scale (Luzon). Lastly, there was no watershed value table grouping, giving one RF model per watershed or simply no regionalization. Watershed-level streamflow modelling is very common in poorly gauged watersheds using process-based methods and these watersheds are partly categorized as PUB (Loukas & Vasiliades, 2014). We wanted to check how streamflow data gaps can be compensated with and without regionalization.

##### Clustering by PCA

As mentioned, the PCA-clustered method relied the watershed grouping based on principal components in light of finding homogeneous watershed clusters (Kahya et al., 2008). This method “normalizes” the watersheds based on the least possible variability among the watershed value tables (also known as dimension reduction). From this, PCs are constructed and interpreted on a dimensionless planar coordinate system with *x* and *y* axis as the first and second PCs (PC1 and PC2) - the components that captures most information of the watersheds. Inside this “PC graph” are the normalized watersheds, plotted and labelled by watershed names. From that, watersheds are clustered using *K-means* algorithm (Hartigan & Wong, 1979), an unsupervised and distance-based clustering method. Clustering depends on how close the watersheds are to the cluster centroid in the PC graph. Centroids are assigned at locations with the least variability among surrounding watersheds. The gap between the centroid and the watershed is measured by a straight line (Euclidean distances) in multiple iterations. The confidence of the clustering is measured when within-cluster variation is minimized, which is measured by the sum of squares (SS) computed by :

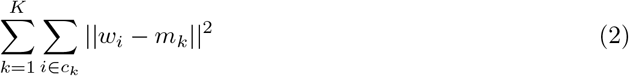

where at the PC graph, centroids are *m*_1_, *m*_2_, … *m_k_*, watersheds are *w_i_* = (*w*_*i*1_, … *w_ip_*), and clusters are *c*_1_, *c*_2_, … *c_k_*.

We first tested the optimal number of clusters, measured by average “silhouettes”, to evaluate the distinction power per cluster number (Rousseeuw, 1987). Out of five possible clusters (2 to 6), we chose the median value (cluster=4) to consider both cluster number (n) and distinction power (silhouette value tipping point). This clustering explained 80% of the dataset variability, computed by dividing between cluster SS from total SS. The importance of the clustering to each observation (watersheds) is measured by cosine squared (*cos2*) (Abdi & Williams, 2010).

The watershed clusters and other key information per watershed are shown in Table 2. The clustering, in general, can be attributed to the homogeneity from general watershed characteristics - size, climate, main land covers and streamflow information.

**Table 2.** Cluster assignments of watersheds along with other key watershed descriptor and information. Climate types include type 1 (distinct wet and dry), type 3 (less pronounced wet and dry), and type 4 (rainfall is more or less distributed yearly). The water (regulating) infrastructures consist of dams, water impounding structures, small reservoirs, and so on.

| Watershed | PCA cluster | Size (ha) | Dominant climate | Cropland, forest, grassland proportions (%) | Mean stream-flow ($\text{m}^3/\text{s}$) | Water infrastructures (number) |
| --- | --- | --- | --- | --- | --- | --- |
| aarb_a | 1 | 236,119 | Type 3 | 5, 93, 2 | 228 | 3 |
| aarb_n | 1 | 19,277 | Type 3 | 0, 99, 1 | 17 | 0 |
| abrb_s | 1 | 20,143 | Type 1 | 17, 81, 1 | 3 | 0 |
| arb_b | 1 | 31,609 | Type 1 | 35, 44, 1 | 3 | 0 |
| arb_c | 1 | 36,507 | Type 1 | 23, 74, 2 | 19 | 0 |
| crb_a | 1 | 104,525 | Type 4 | 24, 70, 6 | 65 | 2 |
| crb_be | 3 | 27,211 | Type 3 | 13, 84, 2 | 3 | 4 |
| crb_bu | 4 | 1,851,853 | Type 3, 4 | 30, 63, 1 | 784 | 15 |
| crb_d | 3 | 169,590 | Type 3, 4 | 6, 92, 1 | 53 | 0 |
| crb_j | 1 | 293,539 | Type 4 | 15, 84, 2 | 210 | 1 |
| crb_m | 3 | 215,601 | Type 1 | 11, 87, 1 | 127 | 15 |
| crb_p | 1 | 317,514 | Type 4 | 12, 85, 2 | 415 | 0 |
| crb_s | 1 | 85,267 | Type 3 | 2, 97, 1 | 45 | 2 |
| crb_t | 1 | 66,994 | Type 4 | 10, 75, 4 | 52 | 5 |
| crb_u | 4 | 1,187,381 | Type 3, 4 | 33, 61, 6 | 454 | 15 |
| mrb_s | 3 | 63,445 | Type 1 | 14, 82, 3 | 34 | 1 |
| prb_a | 2 | 622,280 | Type 1, 3 | 51, 43, 2 | 281 | 7 |
| prb_b | 1 | 37,406 | Type 1, 3 | 82, 12, 6 | 9 | 1 |
| prb_c | 1 | 84,287 | Type 3, 4 | 15, 83, 2 | 78 | 0 |
| prb_p | 1 | 85,739 | Type 1 | 42, 54, 4 | 120 | 0 |
| prb_r | 1 | 147,804 | Type 1 | 72, 25, 3 | 47 | 1 |

**Table 3.**
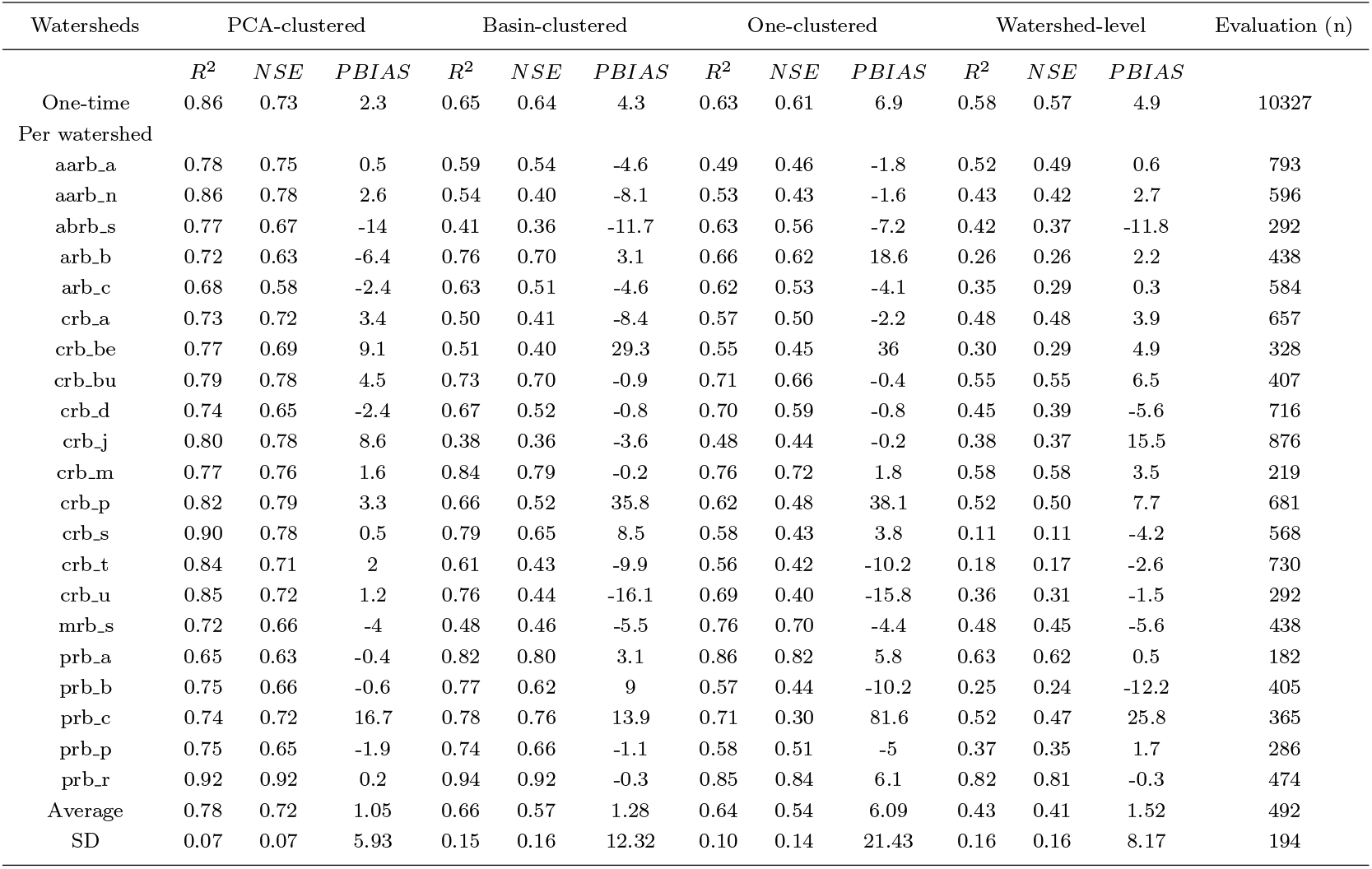
Model evaluation results of the four regionalization methods showing results of goodness-of-fit measures at watershed-level and that of one-time evaluation that used collated the watershed-level evaluation data.

### 3.3 Models evaluation

As in other regression model tasks, we assessed which among the four methods leads to better RF model regionalization by evaluating the RF models per watershed. To evaluate, we used a randomly held-out 20% out of total observed streamflow data from each of the 21 watershed for comparison with the predicted streamflow. We perceived this proportion to be sufficient in retaining the integrity of unseen data given the high variability in our study area and a long observed streamflow time series from 2000 to 2016. Also, ideal data on other local streamflow stations in the Philippines are unavailable yet. We performed per watershed evaluation and a one-time evaluation using each held-out and the accumulated held-out watershed data, respectively. This two-way evaluation provides inter-method and inter-watershed comparison.

Three accuracy metrics were used for evaluation, reported in summary tables. First is the coefficient of determination or *R*^2^ (equation 3) computed as 1 minus the squared difference between the observed *y_i_* and predicted streamflow 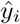; divided by the squared difference of *y_i_* and the mean observed streamflow 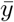. Second, we used Nash Sutcliffe Efficiency or *NSE* (equation 4), computed similarly to *R*^2^ - but instead of *y_i_* - denotes the predictions as *y_i,sim_*. The former is a goodness-of-fit measure of a regression model since we used RF for regression, while the latter measures how well a “model simulation” can predict streamflow relative to 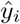. We used these relative coefficients to make results comparable to all watersheds. Though (H. V. Gupta et al., 2009) suggested caution in using *NSE* to evaluate streamflow models in highly seasonal areas, the high spatio-temporal properties of the watersheds (this study) and randomness in selecting evaluation data could compensate for that (H. V. Gupta et al., 1998). Lastly, we used Percent Bias (*PBIAS*) to give an overview whether the RF models over or underestimate the predicted streamflow. *PBIAS* should be 0.0 to be optimal and is computed in equation 5 using the previously mentioned components.

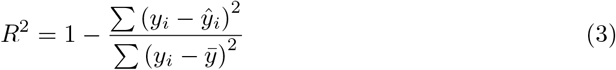

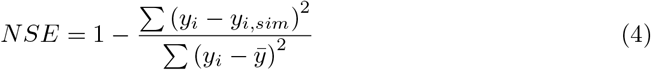

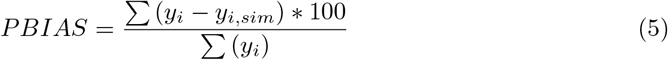

The regionalization methods were also evaluated relative to the major river basins and other categorical watershed characteristics. The former is useful in the context of river basin management in the Philippines, while the latter attempts to find physical and climatic causality from the results.

### 3.4 Variable importance

The main advantage of RF as a machine learning algorithm is to give an importance value to the covariates. RF provides variable important measure (VIM) as a model output, measured by mean decrease in accuracy and equivalent to the “predictive power” of a covariate. This works by permuting a covariate of interest while the rest have their original values. This gives a new prediction in the end and compared with that of the original (no-permutation) model using RF’s test (out-of-bag) data or 1/3 of the total watershed value table. The comparison gives an average error rate difference with and without permutation. The higher the value, the more the covariate influences the model (Breiman, 2001).

VIM was assessed separately for static and dynamic covariates since permutation can be more sensitive to dynamic variables (Altmann et al., 2010). This could overshadow the added-value of static covariates in RF model regionalization. As such, the VIM of static and dynamic covariates were summarized both in absolute values per covariate and % breakdown per covariate category (e.g. season, land cover, slope, elevation).

### 3.5 Uncertainty assessment

Uncertainty is a range of possible predicted streamflow where the true streamflow lies and therefore should be accounted. In this study, we reported uncertainty through confidence intervals relative to a random forest tree. Recall that a streamflow prediction is the mean of all tree predictions in a random forest. To estimate CI, we used the infinitesimal jackknife (IJ) modified by (Wager et al., 2014) and integrated in the *ranger* package. The modified *IJ* approach deals with the uncertainty influenced by the covariates redundancy and limit (*split variable*) during tree growing, and an inherited noise when bootstrapping the watershed value tables - both cannot be dealt by tuning the RF hyperparameters e.g. split variable alone. This is relevant to our covariates of choice because those with yearly values are duplicated more than a daily value. The modified *IJ* is written as:

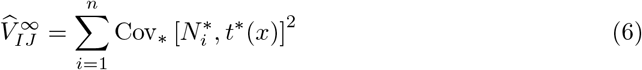

where the variance of a random tree 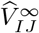 is a function of the covariance 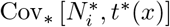 between the random tree *t* * (*x*) and 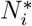 covariate multiple entries in a bootstrap sample the *i^th^* times. To derive the CI at 95% denoted as *CI*^95^, the predicted streamflow 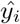 is added or subtracted to the product of 1.96 and the 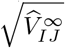 as shown in equation 7. We also derived the percent coefficient of variation (CV%) as an uncertainty summary statistic per regionalization method, where the 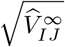 is divided by the mean predicted streamflow 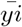 multiplied by 100 (equation 8).

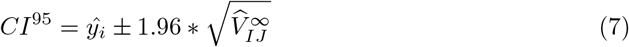

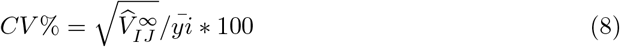

After estimating *CI*^95^, we further analyzed the comparison of observed and predicted streamflow. Using one-time evaluation results, we graphed the comparison relative to a 1:1 line and zoomed into error-prone streamflow ranges. To further analyze, we made a hydrograph of a certain year to spot seasonal errors that are systematic (bias). To highlight seasonal bias, the comparison was aggregated into monthly flows to partially remove random errors (Hamilton & Moore, 2012). Per month and per regionalization method, standardized residuals (SR) were derived to make seasonal results comparable. SR is derived by dividing the residuals 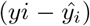 over the uncertainty 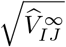. Negative values indicate streamflow overestimation while positive values denote underestimation. We then created figures of monthly SR per regionalization method in a line graph and per watershed in box plots where the lower and upper hinges correspond to the 25^th^ and 75^th^ quartiles. Aside from seasonal bias, we also looked at bias on extreme flows using 50 highest flows of the evaluation data.

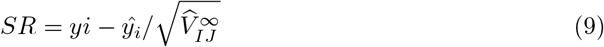

## 4 Results and discussion

### 4.1 Evaluating the regionalization methods

Overall, the observed and predicted streamflow were correlated in all regionalization methods; but the goodness-of-fit indicators varied per method. Results of one-time evaluation indicated that the PCA-clustered method had the highest accuracy, with at most 0.35 and 0.31 higher *R*^2^ and *NSE*; and at most 5% less bias than other methods. Results of basin-clustered and one-clustered methods were identical while watershed-level method was the least accurate. Same outcome was observed in watershed-level evaluation, where the PCA-clustered method showed the highest accuracy in majority (14 of 21) of the watersheds. This method also had the lowest standard deviation (SD) from the three accuracy metrics, suggesting consistency in per-watershed results. Notably, higher accuracy margin was observed in 6 of the mentioned 14 watersheds, outperforming the other models e.g. by 0.25 *R*^2^, on the average. In contrast, the watershed-level method was the least accurate in 20 out of 21 watersheds and with the highest results’ SD. Per watershed results of basin-clustered and one-clustered methods were similar on the average, but varied per watershed. While there was no watershed from the PCA-clustered below the significant thresholds for *R*^2^ and *NSE* (0.6 and 0.5, respectively), there were rare cases that PCA-clustered had high *PBIAS* e.g. *abrb_s* and *arb_b*.

Results slightly varied when analyzed in the context and scale of major river basins. The PCA-clustered method favored basins *aarb* and *crb*, e.g. having 0.24 and 0.15 higher *R*^2^ than the second-best method. Watersheds inside the latter basin also had the most improvement, with 0.49 and 0.26 increase in *R*^2^ and *NSE*, on the average, when using PCA method. Interestingly, the basin-clustered was on par with PCA-clustered for *arb* and *prb*, just 0.02 less *R*^2^ for both basins. Similarly, the PCA-based method was identical with one-clustered in cases of *mrb* and *abrb*.

The results were also analyzed according to climate type, watershed size, and watershed clusters (see Table 2). In terms of climate, the PCA method was favorable for all the climate types especially those under type 4 climate (distributed rainfall, year) with 0.30 *R*^2^ more than the second-best method. In terms of size, the PCA-clustered method was better for relatively small and large watersheds with 0.13 and 0.09 *R*^2^ more than the second-best. In terms of the watershed clusters, it favored its own method being at least 0.10 *R*^2^ ahead from the next-best method. Defying the common finding was watershed *prb_c* in cluster 3, favoring the basin-clustered method.

More generally, the accuracy results seemed to be independent to the number of training (and evaluation) data. The two were uncorrelated at only 0.01 *R* when regressed. On a different note, despite the similarity of *R*^2^ to *NSE*, there were watersheds with higher gaps between the two metrics. Examples are *abrb_s* and *crb_u* with at least 0.05 *R*^2^*-NSE* gap consistent among the four methods. The first happens to be smallest watershed; while the second is not only one of the largest, but also a lone cluster. This suggests that *NSE* reacts better to inter-watershed variability than *R*^2^.

### 4.2 Watershed clustering effect

The PCA-clustered method that outperformed the other methods can lead to the following realizations: (1) homogeneously clustered watersheds can lead to better RF model regionalization, (2) high watershed variability at major river basins and Luzon scales, and (3) watershed-level method is often inadvisable in data-driven methods.

In discussing the first observation, the watershed clustering scheme should be linked to the watershed characteristics. Figure 5 shows how the covariates correlate with the PCs and with themselves in a 2d space (also called as contributions). These confirm the 80% variability explained (in section 4.2) by land cover and physical covariates during the clustering, as they were correlated with the PCs and clumped in the contributions graph. This suggests that the watershed clustering is more of a biophysical representation. The review by Beck et al., (2016) about the history of regionalization highlighted that regionalizing based on similar attributes are effective and even more effective when using single models. Related findings in literature both affirmed that watershed physical features like slope, elevation, and forest cover greatly influence a PC-based watershed clustering scheme (Chiang et al., 2002; Razavi & Coulibaly, 2013). More complex methods combine physical and streamflow signatures to cluster watersheds e.g. (Boscarello et al., 2016). Since streamflow signatures were not included in our clustering due to data limitation, the relevance of streamflow in watershed clustering is confined with the high correlation of daily streamflow (*O.obs*) to principal components (see Figure 5 again). Nevertheless, knowing that streamflow is linked with the biophysical data is a good sign in the PUB context. Ungauged watersheds could use regionalized RF models e.g. PCA-clustered 1 assuming that ungauged watershed is assigned to cluster 1. That would only require ungauged watersheds to have consistent covariates like the 55 covariates (this study) - all from open data.

**Figure 5.**
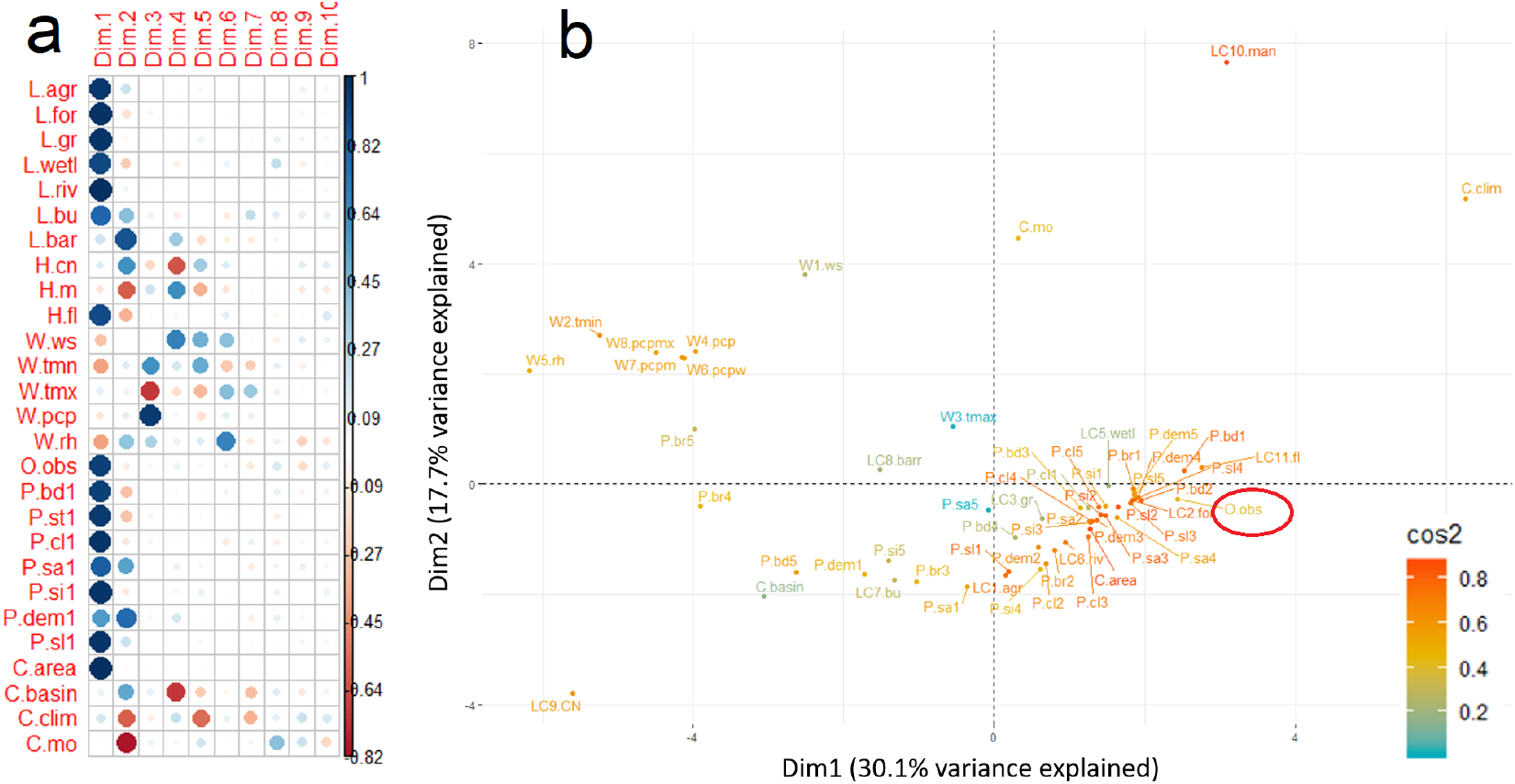
**(a)** Collinearity of the covariates to the PCs - also labelled as *Dim* - showing how physical covariates (*P* and *L* labels) correlate with the main PCs *Dim1* and *Dim2*. It shows one class per physical covariate because results are similar among the 1-5 classes; **(b)** Collinearity of all covariates at 2d space of *Dim1* and *Dim2* showing how physical covariates are clumped, and along with streamflow (*O.obs, encircled*). Clumped covariates influence the watershed clustering the most. High *cos2* orange colors means better agreement of PCs to the covariate locations.

A homogeneous cluster out of watershed physical features can result in two things. One is the imbalance in the number of watersheds per clusters; second is cross-basin grouping where two watersheds apart and belonging to separate basins can be grouped. For instance, PC-based clustering by (Chiang et al., 2002) resulted into a cluster with 59 watersheds and a cluster with one watershed. Similarly (Razavi & Coulibaly, 2013) had a cluster of 37 and another with seven. We can relate from these examples as we have two low-count clusters. In particular, cluster 4 has two watersheds which are both forested; and both with high streamflow and regulated flows. Another cluster only had one watershed, the only large agricultural watershed from the study area. This further suggests that a neighbor-based clustering (spatial proximity) would not be the first option in our study area. On the other hand, we also noticed some flaws from the PCA-clustered method. Two watersheds, *abrb_s* and *arb_b*, had relatively high *PBIAS* despite high *R*^2^ and *NSE*. These are two of the smallest and highly seasonal watersheds mixed in cluster 1. If another cluster is to be made (cluster=5), the two could be isolated. This shows that an optimal watershed clustering involves modelling choices, which should follow best-practices e.g. parameter tuning and iterations especially when upscaling the study.

For the second observation, we assert that homogeneity is not assured at basin-level as watershed characteristics and response can abruptly change in space (He et al., 2011). This is echoed from our results where the basin-clustered method was 0.21 less *R*^2^ than the PCA-clustered method and either second or third best model in basin-level results. This can either be attributed to lack of watershed representation in basins or gaps in time series data (e.g. *arb, abrb, aarb*, and *mrb*). Nevertheless, limitations on data gaps and representation are not evident in basin *crb* - the largest, best represented, and densely gauged basin - which performed best in PCA-based model. This suggests that *crb* can be the most variable basin and this variability is worsen by the presence of water-regulating structures (see Table 2). An exception is basin *prb*, an agricultural basin with less feature variability and fewer water-regulating structures. Similar to the basin-clustered model, the results of the one-clustered model hints that even a lumped watershed information can be insufficient to predict streamflow accurately. Logically, this is the case where watershed variability is at most because of ungrouped and mixed watershed information. In relation, we found that the number of training (and evaluation) data was a non-factor in predicting streamflow accurately. This implies the complexity in “learning” the hydrologic system, where big data could still be insufficient if there is no watershed clustering. Though there are seldom cases where one-clustered method was the best regionalization method e.g. *prb_a* and basin mrb. But by consistency measures, the method choice can still go to the PCA-clustered method.

That is more advisable when the choices include the watershed-level method. The main reasons why watershed-level was the least accurate is due to data scarcity and regionalization itself. watershed-level methods produce stand-alone watershed models trained from a single watershed value table with data gaps. This makes an independent model and literally non-regional, and we say more ideal in process-based and less ideal in data-driven methods. In data-driven methods though, gaps in observed data can be compensated because of watershed regionalization.

### 4.3 Important covariates

In data-scarce basins, the effect of regionalization, regardless of the grouping scheme, can have added value. Information from other watersheds benefited the PCA-clustered, basin-clustered, and one-clustered methods, being way more accurate than the watershed-level method. This can be reflected by the Variable Importance Metric (VIM) given by RF models, which indicates the predictive power of a covariate the way it affects model accuracy once permuted. In watershed-level models, static covariates had 0 VIM values logically because they are static or singled-value - and so in bootstrapped samples - making permutation nonsensical. In other words, 75% of total covariates are of no-use in model prediction. But when watershed information are grouped, the VIM values of static covariates increased from 0 to 22% VIM e.g. in a lumped grouping of one-clustered method. From this, Figure 6 shows both absolute and relative VIM values for static and dynamic covariates. The relative value (% breakdown) is the summary of VIM average per covariate category.

**Figure 6.**
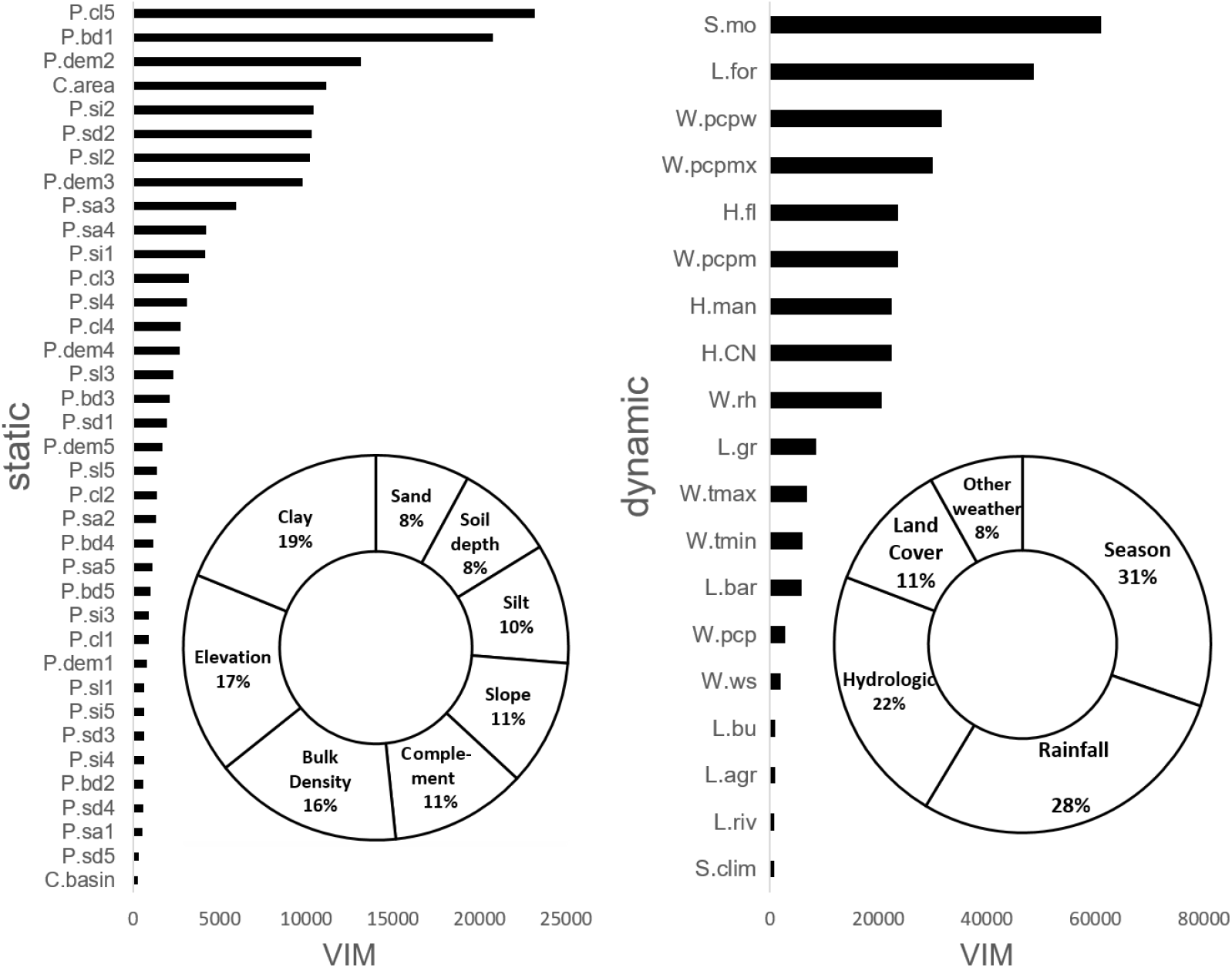
Variable importance measure of static (left) and dynamic (right) covariates of the one-clustered method where the most watershed information are. The bar graphs depict the absolute VIM value per covariate while the pie-charts are summary of % breakdown of VIM average values (uni per covariate category.

The topmost static covariates based on VIM were bulk density, clay, elevation, sand, silt, and slope. VIM appears to be sensitive on dominant covariates (Altmann et al., 2010), which complements the watershed grouping scheme out of physical variables. In particular, very low bulk density (*p.bd1*) and very high clay soil (*P.cl5*) had the highest VIM scores. High clay and low bulk density are naturally coinciding (S. C. Gupta & Larson, 1979) and these characterize most headwaters of our study area. This further reflects that the RF model could relate to some theoretical impacts of static covariates to streamflow. For example, clay soils can hold more water and more regulated run-off (Greacen & Sands, 1980) and watersheds in high altitudes have more rain (Winiger et al., 2005). This is highlighted in Figure 3 in a decision tree context. These realities can also be analyzed in parallel with land cover change. Moreover, sensitivity analysis and co-dependence of covariates to streamflow can give a shed of causality in interpreting RF models (Borgonovo et al., 2017) (Greenwell, 2017).

The VIM of the dynamic covariates were dominated not only by forest land cover class but also by seasonal and weather covariates, contributing 66% of the VIM total. This suggests that RF models are highly sensitive to seasonal weather patterns in the region. Same is observed with other RF models of (Yang et al., 2016), (Reynolds & Shafroth, 2016) where the most dynamic variables were also the most sensitive to VIM. This seems logical given that daily weather data, for instance, have wider range of permutation values than yearly data. Higher temporal resolution is likely the reason why dynamic covariates have higher VIM values than static covariates. For instance, forest cover (*L.for*) became dynamic due to net changes from yearly forest loss, thus increasing its VIM. High forest cover (and changes) of the study area in the past 16 years may have resulted this model sensitivity. As mentioned, land cover change and forest loss effect to streamflow would be interesting to analyze next.

High RF model sensitivity to season and weather covariates can be reflected also during watershed clustering. These covariates dominated the third PC while other dynamic variables appeared until PC 6. This shows how uncorrelated these covariates are from the PCs and the most useful if data-trimming will take place (Chiang et al., 2002). However, in our case, PCA was used for clustering and not for data reduction. The potential overfitting out of correlated covariates are dealt when the split variable and number of trees of RF were tuned to an optimal value. Moreover, random forest automatically de-correlates trees at the most, being lesser prone to multicollinearity (Breiman, 2001).

One key message of this section is the added-value from the static covariates after watershed grouping. These data inputs are all open data, complementing the data-driven method to model streamflow. Moreover, these inputs are relatively very high resolution and therefore advantageous than coarser inputs e.g. the 250m soil inputs used in this study was found to capture soil dynamics better in streamflow modelling (Duku et al., 2015). The advantages from input data can lead into cross-region and even cross-country regionalization.

Worth emphasizing also are the advantages in using random forest via *ranger* package. Aside from giving importance values, another advantage when using RF is to estimate the uncertainties in predicted streamflow.

### 4.4 Uncertainty of random forests and seasonal bias

Despite tuning RF hyperparameters, there remains risks from biased bootstrapping and tree growing in using dense (and correlated) watershed value tables to create RF models. For accountability, we estimated uncertainties in predicted daily streamflow using the modified *IJ* approach.

Error assessment is an important post-modelling steps in streamflow modelling (Abbaspour et al., 2017). Shown in Figure 7 are the results of one-time evaluation of the four regionalization methods with the following sub-figures: (a) predicted streamflow and its confidence intervals CI^95^ aggregated to calculate the CV%; (b) zoomed-in results until 1000 *m^3^/s*; and (c) evaluation result in a sample year (2006, all watersheds data) to highlight over and under estimation.

**Figure 7.**
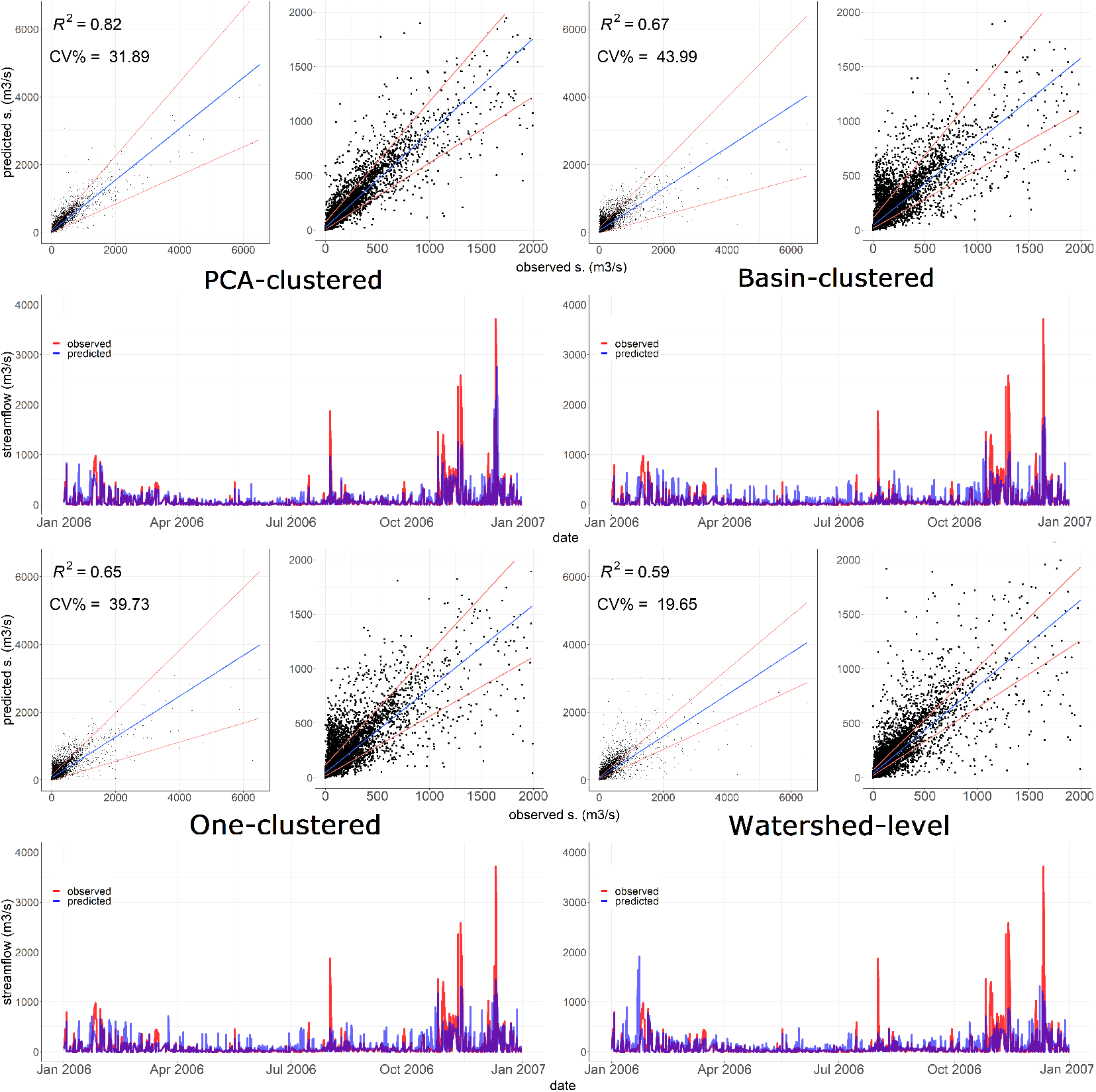
Evaluation results of the four regionalization methods using the accumulated 20% held-out data of watersheds to highlight uncertainty and errors: Upper left graph (a) shows the all predicted and observed streamflow comparison with CI^95^ range, and the R^2^ and CV%. Upper right graph (b) zooms to below 1000 *m*^3^/*s* flows to highlight the systematic trend or bias. Lower graph (c) shows time-series of the observed and predicted streamflow in year 2006. The full time series hydrograph is shown in the appendices.

In Figure 7(a), watershed-level method had the least uncertainty (13% less at least) among others, followed by the PCA-clustered, one-clustered, and basin-clustered methods. This is potentially caused by the trade-off between grouping and not grouping watershed information. The CI are computed to account for the minimized noises and bias of the bootstrapping procedures (Wager et al., 2014). Given that watershed-level value table for model training is unmixed with other watersheds, it can have less variable random subsets. Same logic is applicable on why the clustered (more homogeneous) was the least uncertain among models with watershed grouping. Another noticeable result was the higher uncertainty from the highest or extreme flows. These flows are outcomes of heavy rainfall e.g. typhoon season from July to October, occurring several times in a year (Cinco et al., 2018). Being the minority among the flows can result into higher uncertainty, related to streamflow data imbalance or histogram skewness (Bhattacharyya, 2013). This is quite common in streamflow data especially in areas with very pronounced season. In cases where certain predictions of extreme flows are needed for specific applications like flood modelling, observed streamflow should adhere to sampling protocols (Tian et al., 2013) and prefer data with higher temporal resolution e.g. hourly (Saadi et al., 2019).

Also observable from the figures are systematic over and under estimation of streamflow (bias). Using averages of the 50 highest flows, we derived underestimation of extreme flows to be 53%, 113%, 120%, and 127% from PCA-clustered, basin-clustered, one-clustered, and watershed-level methods, respectively (see full hydrograph in Appendix B). While these flows seem to be the most uncertain and underestimated, as mentioned, these are minority flows. For general streamflow modelling applications (e.g. water resources accounting) unlike flood mitigation, extreme flows can be averaged out. The more prominent error source was the overestimation until around 1000 *m*^3^/*s*, as highlighted in Figure 7(b). The correction of these relatively lower flows was more evident in PCA-clustered method. In a sample year in Figure 7(c) (full time series in Appendix B), reduced overestimation of low flows and reduced underestimation of high and extreme flows were also evident in PCA-clustered. Nevertheless, analysis of bias gets clearer when looking at aggregated flows.

The predicted streamflow was analyzed per month to understand bias related to seasonal patterns. This analysis per regionalization method is shown in Figure 8, while the results per watershed are shown in Appendix A. Visualizing monthly streamflow is also strategic in the sense that the RF models were very sensitive to month (*C.mo*) covariate. Using the monthly flow aggregates, we first compared the standardized residuals, calculated in equation 9. In general, the graph depicts streamflow overestimation in all models and most months. Similar to the evaluation results, the clustered model was the least bias in 8 out of 12 months.

**Figure 8.**
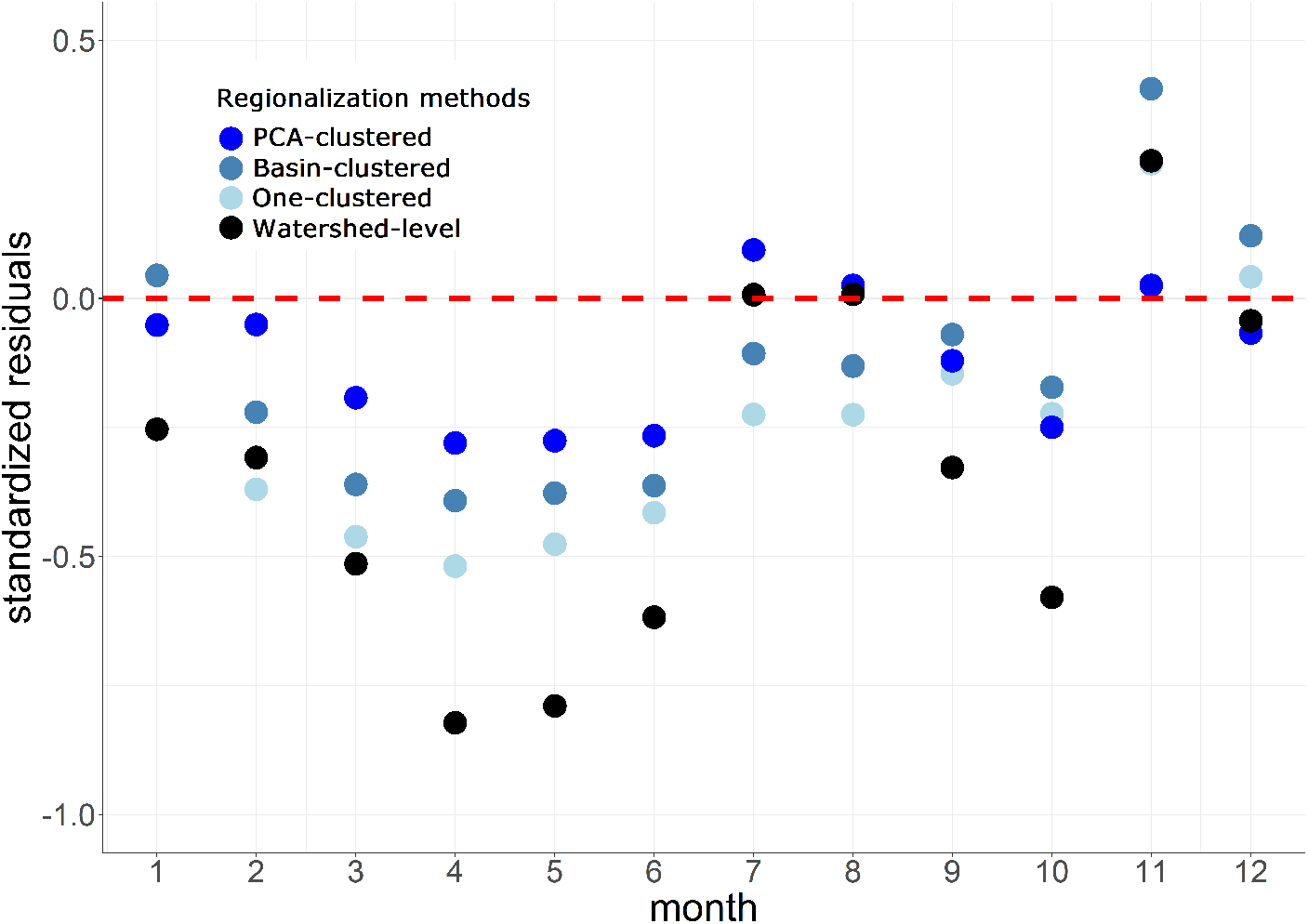
Standardized residuals (*observed* - *predicted*) / uncertainty of monthly average flows to show trend of seasonal errors and its magnitude. Below the 0 line depicts that predicted streamflow is overestimated and otherwise if above the line.

Seasonally, the monthly bias graph depicts three trends common to the four regionalization methods: (1) overestimation during the dry season of March until June, (2) brief underestimation during monsoon season from November to January, (3) lower bias in wettest months of July to September. In these cases, streamflow predictions can be reactive to seasonal weather patterns (Singh, 1997). On the other hand, streamflow models can also capture non-linear watershed behaviors. For instance, predicted streamflow did not increase despite heavy rains because of water storage effect in the landscape (Shortridge et al., 2016). In theory, streamflow should increase after rainfall events. However, in times of high streamflows even without rainfall, streamflow can be influenced by both natural and man-made water regulations.

The first observation can be linked to the effect of water-regulating structures, mainly dams for irrigation, which release water particularly during the first cropping season (1^st^-2^nd^ quarter) and vary in water release schedules. These structures often exist in watersheds with relatively large lowland agriculture e.g. aarb_a, *crb_bu, crb_m, crb_u*, and prYa. Despite having higher errors in dry months, these watersheds benefited from watershed clustering and performed poorest using watershed-level model. A watershed-level model can be more inadvisable when watersheds have water-regulating structures by having no extra information to compensate for irregular flow time-series and data gaps. In any future attempts to minimize dam effects, streamflow signatures from flow duration curves could be included in watershed clustering as well as dam-related data as covariates (assuming data availability).

For the second observation, after the wettest months, the aquifers and shallow groundwater becomes full, thus retaining relatively higher streamflow even without rains. Moreover, the study area is affected of northeast monsoon making December to February the coolest months. Soil moisture evaporation is less pronounced in lower temperatures thus retaining high streamflow (Penman, 1948). This scenario was more pronounced in watersheds from basins aarb and crb - the northeast basins in the study area. Moreover, dams tend to release excess water during this season resulting into above-average streamflows. To model these realities, temporal memory effects and spatial weather inputs could be integrated, resulting into a spatio-temporal streamflow models.

While the first two observations are linked to non-linear realities, the last observation is related to a linear rainfall-runoff scenario, where streamflow is reactive to rain events. In the study area, raining often starts in June and this month was also the tipping point to reverse high errors from dry months. This behavior is common to all watersheds, but more reactive to this seasonal transition are smaller watersheds with minimal regulating structures e.g.abrb_s, arb_b, *arb_c, crb_t, prb_b, prb_r*. Smaller watersheds seem to adhere more in rainfall-runoff events than larger ones. To exemplify, one study used a linear model in a small watershed to forecast drought (Tigkas et al., 2012) while another reported a more pronounced response of smaller watersheds to land cover change (Dias et al., 2015). As mentioned, assessing forest loss effects in different watershed sizes would be an interesting follow-up to this study.

The seasonal bias can also be affected by errors and associated uncertainty from gauged data. While we treated this data as ground-truth, this dataset have been computed indirectly from water depth or stage and curve ratings (JICA, 1998). In a study by (Horner et al., 2018), this kind of measurement can introduce non-negligible systematic errors in addition to negligible man-made errors and negligible non-systematic measurement errors. Accounting this root error source could be difficult given that raw data are often inaccessible. What is more, RF model predictions can be bias to the mean of the dataset resulting into systematic errors, but can be dealt at model-level bias correction methods (Zhang & Lu, 2012).

## 5 Conclusion

We explored regionalizing RF models in mountainous watersheds of the Philippines using biophysical, weather, and other watershed data to represent the main inputs of a hydrological system. These datasets were collated as watershed value tables unique per watershed, and followed four grouping schemes namely PCA-clustered, basin-clustered, one-clustered, and watershed-level - all with distinct RF models to be regionalized. The regionalization methods were evaluated using 20% held-out data of each watershed.

Outcomes of model evaluation amplified that the two-step nature of regionalization (watershed grouping then model transfer) is advisable using our approach. Gains from “outside” watershed information benefited PCA-clustered the most based on goodness-of-fit measures. Results of semi and one-clustered methods suggest that watersheds at major river basin and Luzon scales can be very variable. The watershed-level with no watershed grouping was the least accurate method, as indicated by the insignificance of static covariates and lesser watershed information. These results suggest to cluster watersheds first prior to streamflow model regionalization, preferably using PCs. That would address gaps from observed data in gauged watersheds and observed data absence in ungauged watersheds.

The PC-based watershed clustering is a biophysical representation where 80% of the dataset variability is explained by soil, slope, elevation, and land cover. This coincides with the 22% increase in importance value of physical-static covariates in RF models. On the other hand, covariates weather and month were the most important dynamic covariates, suggesting sensitivity of the RF models to season. The significance of these (mostly open) input data magnifies the increasing momentum on data-driven streamflow modelling. Assuming access to local weather data, our approach can be replicated into other tropical countries. Otherwise, downloadable global weather data can substitute for local data.

Aside from covariates importance, RF (using *ranger* R package) accounts for uncertainty and includes it as an RF model output to measure confidence intervals on predicted streamflow. Such output can efficiently identify those significant predicted streamflow. These RF functionalities are recommended to any RF-based streamflow modelling, and also an in-depth sensitivity analysis of covariates and streamflow to assess their causality e.g. use of partial dependence plots.

Uncertainty from predicted streamflow was minimal (lowest CV%) in watershed-level method among all methods, while the PCA-clustered had lower CV% among those with watershed grouping. This suggests a trade-off between additional watershed information and uncertainty. Most uncertain (wide CI^95^) predictions were extreme flows from typhoon events. Such flows were also underestimated and needs improvement for potential flood modelling uses.

Monthly streamflow from several dry and wet months exhibited overestimation evident to all methods - the PCA-clustered having the least overestimation in 8 of 12 months. That prediction bias conforms to events that cause relatively high streamflow even without rainfall e.g. irrigation water supply in dry season, moist soils and recharged aquifers during monsoons, and post-typhoon days in wet season. These observations further suggest that the streamflow models can be reactive to seasonal and non-linear realities related to water regulation. Therefore, the regionalization method can be fine-tuned to integrate these water regulation effects (e.g by using more data and covariates, integrate “physics”, and switch to deep learning).

Other potential follow-up to this study would be to assess forest loss effect on streamflow and upscale regional PUB nationwide after assigning all ungauged watersheds into clusters.

## Appendix A Standardized residuals per watershed

**Figure A1.**
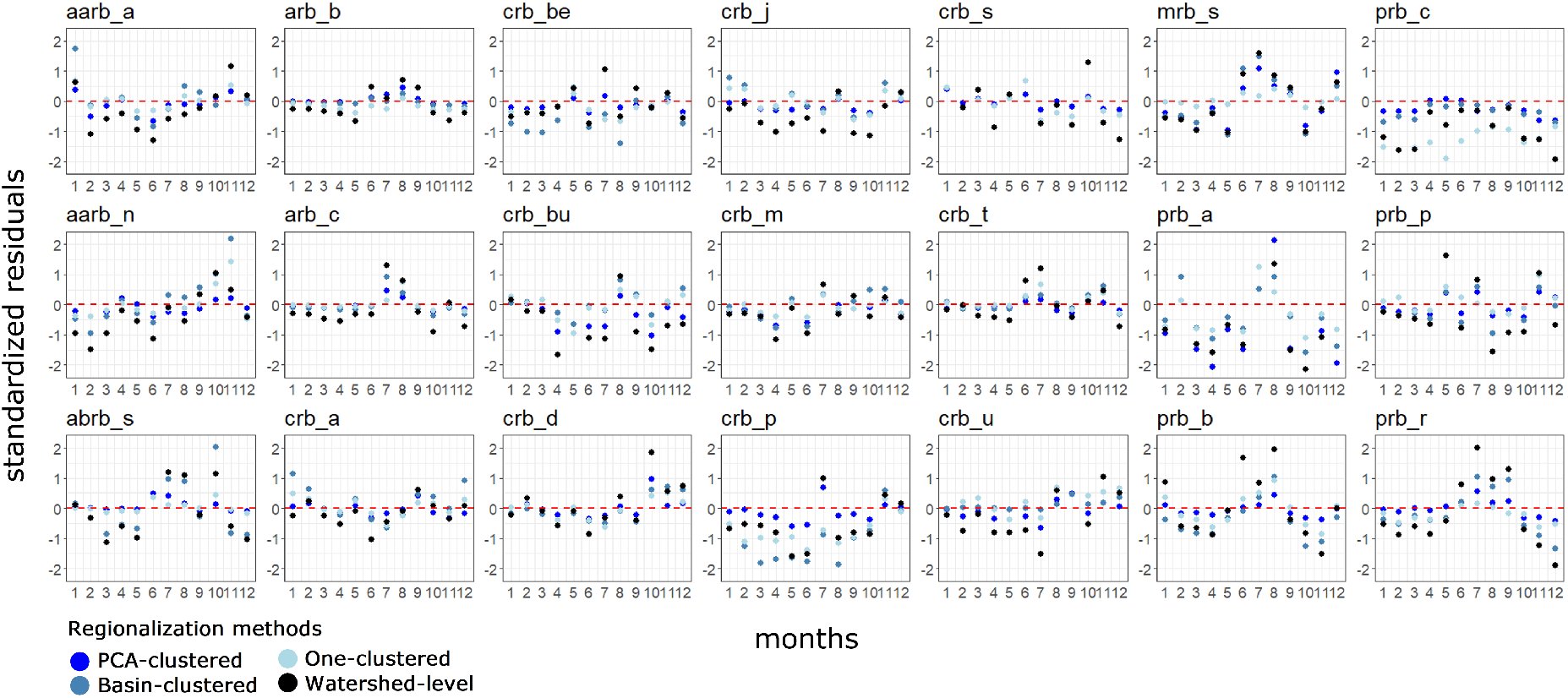
Standardized residuals (observed streamflow - predicted streamflow / uncertainty) per month and per watershed among the four regionalization methods.

## Appendix B Hydrographs of the four regionalization methods

**Figure B1.**
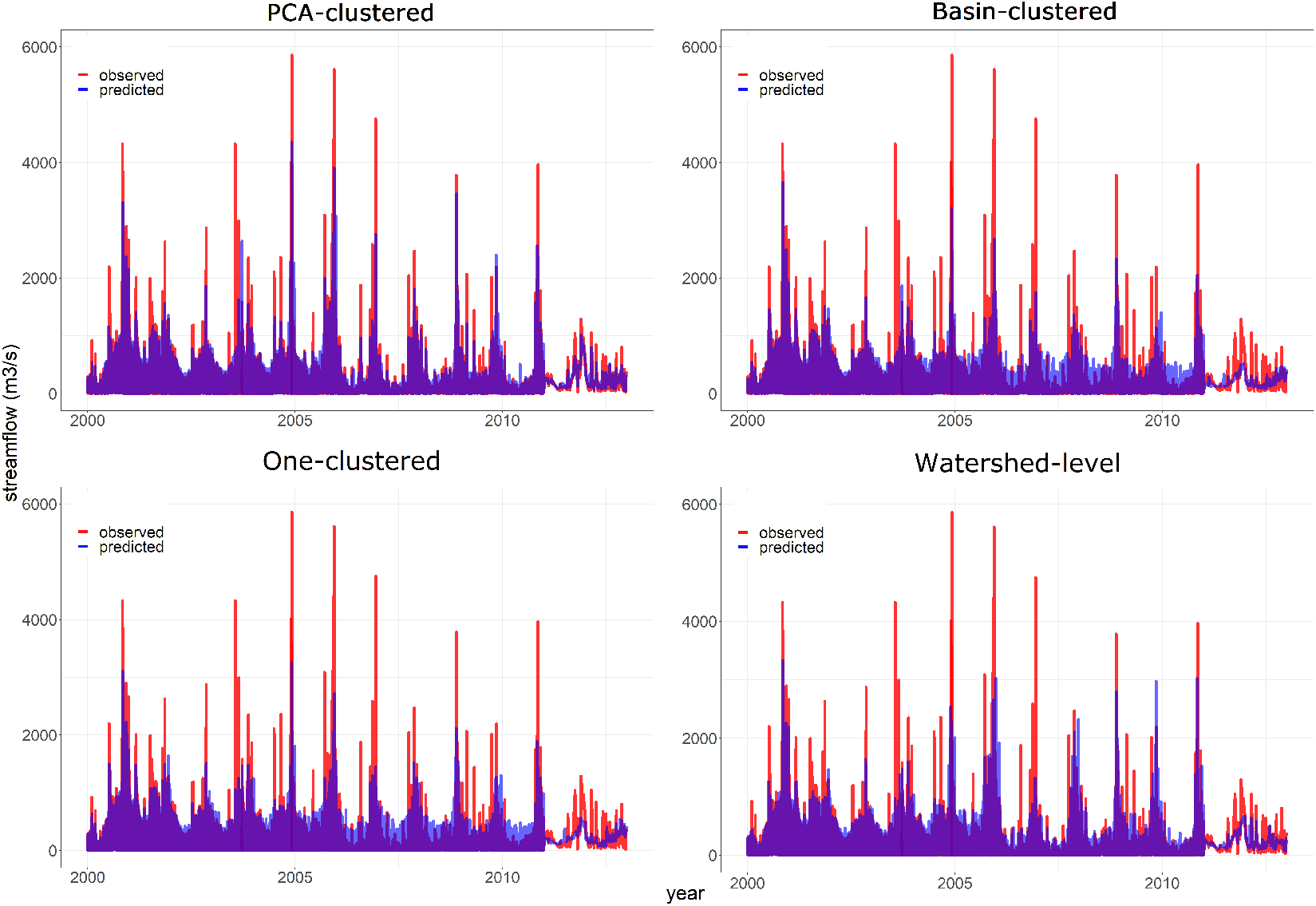
Hydrograph of one-time evaluation showing predicted and observed streamflow, from 2000 to 2016, for the four regionalization methods.

## Acknowledgments

We give credit to key data sources, PAGASA and DPWH, for providing weather and streamflow data from local stations in the Philippines. Special thanks is given to the Forest Foundation Philippines for providing funds to acquire and pre-process the mentioned data. These data should be accessible at http://bagong.pagasa.dost.gov.ph/ for weather and https://apps.dpwh.gov.ph/streams_public/home.aspx for streamflow. Other input data can be accessed online for free as described in the methods section e.g. soil at https://soilgrids.org/#!/?layer=ORCDRC_M_sl2_250m&vector=1. Codes are accessible at https://github.com/arnanaraza/streamflow. We declare no conflict of interest on this research.

